# Screening of MMV open-source libraries using Bunyamwera virus as a model reveals inhibitors of Oropouche virus infection

**DOI:** 10.64898/2026.01.14.699453

**Authors:** Natasha M. Cassani, Hayley M. Pearson, John N. Barr, Mark Harris, Ana Carolina Gomes Jardim

**Affiliations:** School of Molecular and Cellular Biology, Faculty of Biological Sciences, University of Leeds, Leeds, LS2 9JT, United Kingdom; Laboratory of Antiviral Research, Institute of Biomedical Sciences, Federal University of Uberlandia, Uberlandia, Minas Gerais, Brazil

**Author notes:** Corresponding authors: Professor Ana Carolina Gomes Jardim, Institute of Biomedical Science (ICBIM), Federal University of Uberlandia (UFU), Avenida Amazonas, 4C- Room 216, Umuarama, Uberlândia, Minas Gerais, Brazil, CEP: 38405-302, Professor Mark Harris, Faculty of Biological Sciences and Astbury Centre for Structural Molecular Biology, University of Leeds, Leeds, United Kingdom, LS2 9JT.

**Keywords:** Bunyamwera virus, Oropouche virus, high-throughput screening, compound library, antiviral, hit-to-lead

## Abstract

Arboviral infections remain a major public health concern in tropical and subtropical regions, where environmental and socioeconomic conditions facilitate the circulation of diverse RNA viruses, including over 350 members of the recently renamed *Peribunyaviridae* family. In this regard, Oropouche virus (OROV) has caused explosive outbreaks in the Amazon region and shows wider distribution in Brazil, with confirmed neurological infections and fatalities in 2024. The absence of effective antiviral therapies highlights an urgent need for the discovery of anti-OROV drugs. In this work we evaluated the antiviral potential of the Pandemic Response Box and Global Health Priority Box libraries from Medicines for Malaria Venture (MMV) against OROV, using a well-established *Bunyamwera virus* (BUNV) system, widely used as a prototype of the *Peribunyaviridae* family, expressing the reporter gene eGFP. A screening protocol based on fluorescent reporter gene detection by live cell imaging was employed. 9 compounds were identified with significant activity against BUNV-eGFP with 3 being highlighted as the most promising hits: Trimetrexate, GSK-983 and MMV1634385. These compounds were validated by testing for activity against OROV, and in these assays, GSK-983 exhibited the lowest EC_50_ value and consequently the highest selectivity index. Future studies should evaluate the antiviral efficacy of GSK-983 and assess its pharmacokinetic profile. Furthermore, our results indicate that our approach employing BUNV-eGFP as a model against OROV can lead to the discovery of promising and interesting hit compounds, thereby providing a validated screening approach for future antivirals against Oropouche fever.

## 1. Introduction

Arboviral infections remain a major public health concern in tropical and subtropical regions [1]. Environmental and socioeconomic conditions favour the circulation of diverse RNA viruses, including members of the recently renamed *Peribunyaviridae* family (formerly *Bunyaviridae*) [1,2]. This family represents one of the largest groups of negative-sense single-stranded RNA viruses, with over 350 identified members [2,3]. It is named after the Bunyamwera virus (BUNV), which is considered the prototype of the family, and includes five genera, with *Orthobunyavirus* being the largest and most diverse [4].

All the orthobunyaviruses share common structural characteristics: they are enveloped viruses with a genome segmented into three RNA species, designated S (small, ∼700 b), M (medium, ∼4,400 b), and L (large, ∼6,900 b) [5]. These three segments encode four structural proteins, following an expression strategy conserved throughout the family: the S segment encodes the nucleoprotein (N), the M segment encodes two glycoproteins (Gn and Gc), and the L segment encodes the RNA-dependent RNA polymerase (L protein or RdRp) [6]. In addition, most orthobunyaviruses also encode two non-structural proteins: NSs, derived from the S segment, and NSm, derived from the M segment [7,8]. Viral particles are enveloped and predominantly spherical, measuring between 70 and 110 nm in diameter [5,6].

Several orthobunyaviruses have been reported in Brazil, including some that are associated with human and animal diseases [9]. Most of them have been isolated in the Amazon region from mosquitoes and non-human vertebrate hosts, as exemplified by the recent Oropouche virus (OROV) outbreaks [9,10]. OROV has circulated silently in regions inhabited by its vector (*Culicoides paraensis*), however, few studies have addressed its prevalence until 2024 [10–12]. In March 2024, the Pan American Health Organization (PAHO) issued an alert in response to the rapid increase in OROV cases in South America, characterized as an outbreak of the disease with a high risk of infection [11,12]. This outbreak was caused by the upsurge of a new reassorting lineage (OROVBR-2015-2024 or AM0088), genetically distinct from the classical genotypes I-IV. This new reassorted virus had significantly higher titres and a more virulent phenotype, compared to previous strain (OROVBR-2009-2018 or BeAn19991) [12].

Antiviral strategies for orthobunyavirus infection are therefore required to achieve clinical efficacy during outbreaks and address the potential emergence of new reassortants [10]. A large-scale evaluation of previously characterized drugs may identify additional therapeutic options that can be repositioned for accelerated preclinical and clinical evaluation [13].

Here we hypothesized that BUNV, as a prototype *Orthobunyavirus* sharing high genomic and structural similarity with OROV [14,15], can serve as a reliable model for antiviral screening. Therefore, compounds that inhibit BUNV-eGFP replication are expected to display cross-reactive activity against OROV using a BUNV-eGFP as a model. For this, we developed a screening assay, with an automated image-based readout, to identify effective compounds contained within the Pandemic Response Box (PRB) and Global Health Priority Box (GHPB) libraries from the Medicines for Malaria Venture (MMV) against OROV. 640 compounds from both boxes were screened, identifying 9 candidate antiviral compounds, of which 3 (Trimetrexate, GSK-983 and MMV1634385) were identified as promising hits. Of these GSK-983 was also demonstrated to be a promising anti-OROV agent.

## 2. Methods

### 2.1. Compounds

The PRB and GHPB compound libraries were obtained from the Medicines for Malaria Venture (MMV) (Geneva, Switzerland: https://www.mmv.org/mmv-open). Compounds (purity >90%) were supplied as a dry thin film in wells of 96-well plates. They were dissolved in a 10 µL stock solution of 100% dimethyl sulfoxide (DMSO) (v/v) to a final concentration of 10 mM. The stock solution was then diluted to a 1 mM working solution (10 µL of compound diluted in 90 µL of DMSO), and 10 “daughter” plates that were created for the assays containing 10 µL of the working solution to minimize freeze-thaw cycles. All plates were frozen at -20°C in a humidity-protected environment.

### 2.2. Cell culture

Human adenocarcinoma alveolar basal epithelial cells (A549, National Institute for Biological Standards and Control [NIBSC], United Kingdom), baby hamster kidney cells (BHK-21, ATCC #CCL-10) and BSR-T7 cells (derived from BHK21 cells, that constitutively express bacteriophage T7 polymerase) [16] were cultivated in Dulbecco’s modified Eagle’s medium (DMEM; Sigma–Aldrich, Gillingham, United Kingdom) supplemented with 100 U/mL penicillin (Gibco Life Technologies, Thermo-Fisher Scientific, Paisley, United Kingdom), 100 mg/mL streptomycin (Gibco Life Technologies, Thermo-Fisher Scientific, Paisley, United Kingdom), 1% (v/v) non-essential amino acids (Gibco Life Technologies, Thermo-Fisher Scientific, Paisley, United Kingdom) and 10% (v/v) fetal bovine serum (FBS; Hyclone, Logan, UT, United States) at 37°C in a humidified 5% CO_2_ incubator. BSR-T7 cells were cultivated in the presence of Geneticin (G418, Invitrogen, Paisley, United Kingdom) at 1 mg/mL [16].

### 2.3. Rescue of recombinant BUNV harbouring a eGFP reporter

Plasmids containing full-length cDNAs of BUNV genome segments: pT7riboBUNL(+), pT7riboBUNM(+), pT7riboBUNS-eGFP(+) and the T7 polymerase pT7ribo have been described previously [17,18]. A total of 1 μg of each BUNV plasmid and 300 ng of pT7ribo was used to transfect BSR-T7 cells (3 × 10^5^ cells/well in a 6-well plate) using the Lipofectamine 2000 reagent according to the manufacturer’s protocol (Thermo-Fisher Scientific, United Kingdom). After five days, the supernatant was collected, transferred to BHK-21 cells grown in a T25 cm^2^ flask, cells were incubated until complete cell lysis was observed and the supernatant was harvested as P0 stock. Viral titres were determined using plaque assay in A549 cells [19]. Briefly, A549 cells grown in a 24-well culture plate were infected by ten-fold dilutions of virus stock. Following an incubation of 1 h at 37 °C and 5% CO_2_, 0.5 mL of culture medium supplemented with 2% FBS and 2% carboxymethylcellulose sodium salt (CMC) (Sigma-Aldrich) were added, and the incubation was extended for three days at 37 °C 5% CO_2_. After removing the media, cells were fixed with formaldehyde (4%) and stained with 2% crystal violet diluted in 20% ethanol. Plaques were counted and expressed as plaque-forming units per millilitre (PFU/mL) [19]. All the BUNV-eGFP work were conducted in biosafety level 2 laboratories at the University of Leeds.

### 2.4. Wild-type Oropouche virus (OROV)

A wild-type OROV strain AM0088 was kindly provided by Professor José Proença-Modena of the Laboratory of Emerging Viruses of the State University of Campinas [12]. Virus was expanded and titred by plaque assay in A549 cells as described in the section above for BUNV-eGFP.

All OROV infection assays were performed at a BSL-2 laboratory at the Laboratory of Antiviral Research, under the authorization number CBQ: 163/02 and process SEI: 01245.006267/2022–14 from the CTNBio - National Technical Commission for Biosecurity from Brazil.

### 2.5. Cell viability screening assays

To establish the non-cytotoxic concentration of the compounds, a suspension of A549 cells was seeded in 96-well microplates at a concentration of 2 × 10^4^ cells/well and incubated at 37 °C and 5% CO_2_ for 24 h. A screening concentration of 10 µM was used for each compound, diluted in culture medium, and added to the cells in triplicate. DMSO 1% was used as the untreated control. After 24 h, the supernatant was removed, and cells were incubated with a solution containing 3-(4,5-dimethylthiazol-2-yl)-2,5-diphenyltetrazolium bromide (MTT) (Sigma-Aldrich) at 1 mg/ml of culture medium at 37 °C for 30 minutes. The MTT solution was then removed, replaced with the same volume of DMSO, and subjected to absorbance reading at 490 nm, as previously described [20]. The percentage of cell viability was determined by dividing the mean absorbance of the treated wells by the mean absorbance of the control wells and multiplying by 100. Percentages of cell viability greater than 80% were considered as non-cytotoxic [21].

### 2.6. Antiviral assays

To identify the antiviral activity of the compounds, A549 cells were plated in 96-well microplates at a concentration of 2 × 10^4^ cells/well and incubated at 37 °C and 5% CO_2_ for 24 h. Only compounds that showed no cytotoxicity were tested, diluted in culture medium containing BUNV-eGFP at a concentration of 10 µM, with a multiplicity of infection (MOI) of 0.1. Medium containing virus and compound was added to the cells in triplicate. DMSO 1% was used as the untreated control. Plates were placed in the IncuCyte® S3 Live-Cell Analysis system (Sartorius) for 24 h, and green fluorescence light was observed at a 10 × objective. Images were recorded and analysed employing the basic analyser from the IncuCyte S3 system to obtain the average mean intensity of green fluorescence (GCU: Green Calibration Units per well). Values are presented as the mean fluorescent intensity per cell from a minimum of three independent experiments. Hits were identified as compounds that inhibited BUNV replication to a value greater than 1log_10_ reduction (≥ 90%).

### 2.7. Determination of the effective concentration of 50%

Compounds active against BUNV-eGFP were selected for dose–response assays for hit validation through the determination of the effective concentration of 50% (EC_50_). Two-fold serial dilutions of each compound (10 µM to 20 nM) were prepared in infection medium containing BUNV-eGFP (MOI 0.1) and added to A549 cells in 96-well plates. Plates were placed in the IncuCyte S3 system and GCU were measured at 24 hours post-infection (hpi) as described above. Data were normalised according to the equation (T/C) × 100%, where T and C represent the mean GCU of the treated and untreated control groups, respectively.

### 2.8. Mechanistic assays

#### 2.8.1. Early stages

A549 cells at the density of 2 × 10^4^ cells per well were seeded in 96-well plates 24 h prior infection and treatment. In pretreatment assay, cells were treated for 1 h with validated hits prior to the infection, washed with PBS and infected with BUNV-eGFP for 1 h. Then, cells were washed with PBS and fresh medium was added for 24 h. In entry inhibition assay, cells were infected using media containing each compound and BUNV-eGFP for 1 h, washed with PBS and incubated with fresh medium for 24 h. All infections were performed with BUNV-eGFP at an MOI of 0.1 and compounds at 10 µM. Efficiency of virus replication was assessed by measurement of GCU at 24 hpi in the IncuCyte S3 system as described above. DMSO at 1% (v/v) was used as the vehicle control.

#### 2.8.2. Virucidal assay

The virucidal activity was assessed using the same protocol of entry assay, except that inoculum containing compound and virus was incubated for 1 h before it was added to the cells. Efficiency of virus replication was assessed by measurement of GCU 24 hpi in the IncuCyte S3 system as described above. DMSO at 1% (v/v) was used as the vehicle control.

#### 2.8.3. Late stages

In late-stage assays, 2 × 10^4^ A549 cells were infected with BUNV-eGFP (MOI 0.1) for 1 h, washed extensively with PBS to remove unbound virus, and then incubated with medium containing each compound at 2 µM. Plates were placed in the IncuCyte S3 system and GCU were measured at 4, 6, 8, 10, 12, 18 and 24 hpi.

### 2.9. Validation against OROV

Compounds active against BUNV-eGFP were selected for dose–response assays against OROV_AM0088_. Two-fold serial dilutions (10 µM to 20 nM) of each compound were prepared in infection medium containing OROV (40 PFU/well) and added to A549 cells in 24-well plates for 1 h. Then, medium containing CMC at 2% with the same concentration of each compound was added to the cells for 5 days as previously described with modifications [22]. After the treatment, medium was removed, cells were fixed with 4% formaldehyde for 20 min and stained with violet crystal 0.5% for PFU counting. Data were normalised according to the equation (T/C) × 100%, where T and C represent the mean PFU of the treated and untreated control groups, respectively.

### 2.10. Statistical analysis

All data were analysed using GraphPad Prism 8.0 and expressed as mean ± standard deviation. Statistical analysis was first evaluated for Gaussian distribution. Differences were then determined using analysis of variance (one-way or two-way ANOVA) or Student’s t-test for parametric tests, and Mann–Whitney or Kruskal–Wallis tests for nonparametric analyses. EC_50_ values were calculated using non-linear regression with four parameters in variable slope. P values < 0.05 were considered statistically significant.

## 3. Results

### 3.1. Antiviral assay and identification of hits against BUNV-eGFP

To start, we developed a screening assay, with automated image-based readout, to identify effective compounds from the PRB and GHPB libraries (MMV) against OROV. However, given the lack of an available reporter-expressing construct of OROV for rapid screening of compounds, we employed a BUNV system expressing the reporter gene eGFP as a model [17,18]. Initially, assays were performed employing A549 cells to assess compound cytotoxicity by the MTT assay. All compounds were screened at a final concentration of 10 µM.

From the 640 compounds screened for both boxes, 55 compounds presented cytotoxicity (Figure S1), assessed by a cell viability less than 80% [21] and were therefore excluded from the antiviral screening. Then, the 585 compounds were evaluated against BUNV-eGFP employing a live cell-imaging measurement of GCU using the IncuCyte S3 system, which consists of an automated phase-contrast and fluorescence imaging microscope housed in a humidified incubator supplied with 5% CO_2_. This method allows the accurate and precise measurement of viral titres in high/medium-throughput contexts [23]. The minimum practical hit limit for the assay was set at 1log_10_ reduction (≥ 90%). As a result, a total of 9 compounds were selected as hit candidates in the PRB (**Figure 1A**). These compounds showed eGFP fluorescence levels significantly greater than a 1log_10_ reduction compared to the negative control and were selected for validation assays. In the GHPB no compounds were identified as possessing anti-BUNV activity (**Figure 1B**).

**Figure 1.**
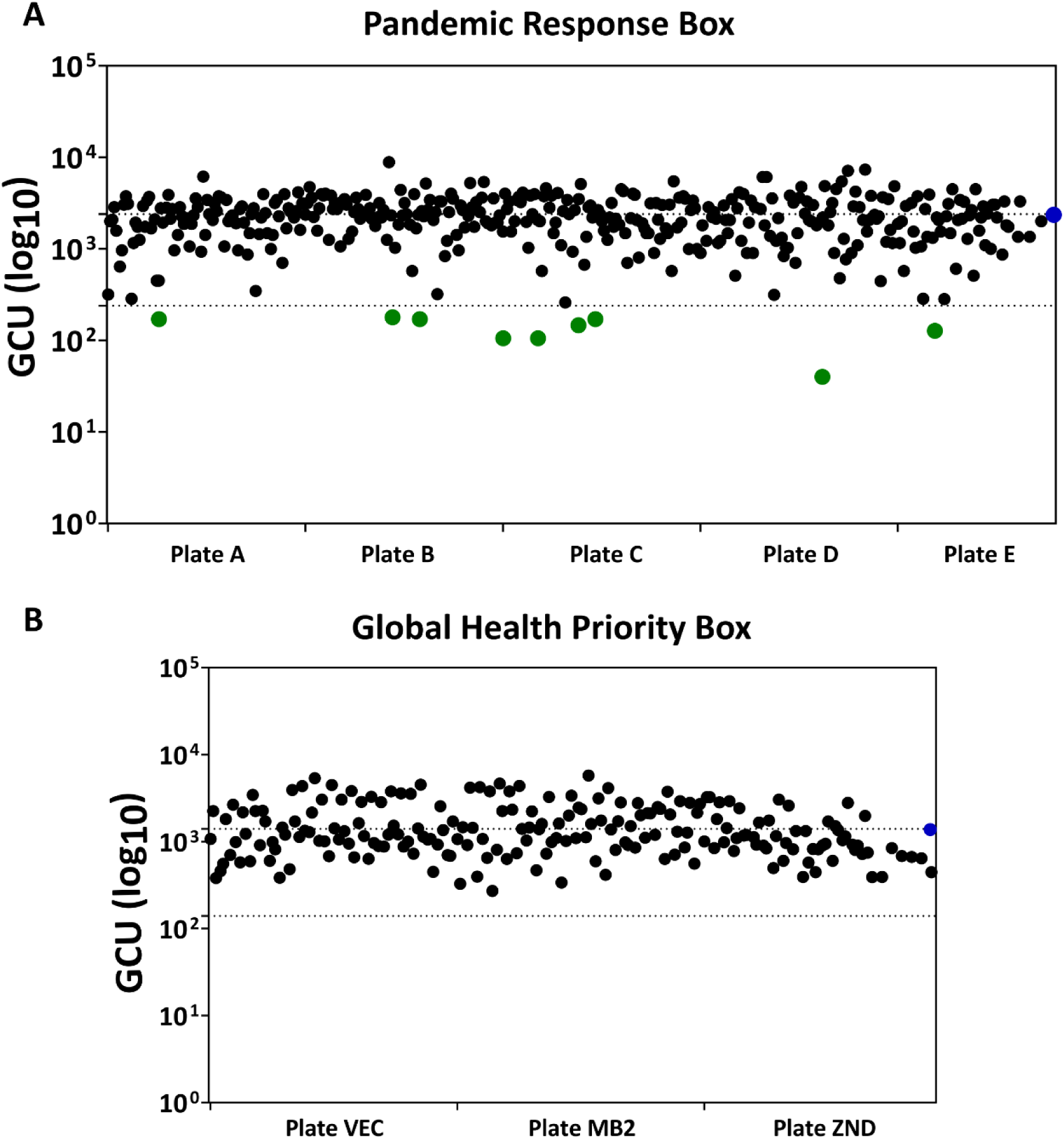
Live-cell imaging MTS for the Pandemic Response Box (PRB) and Global Health Priority Box (GHPB). Scatter plot for screening results from the (A) 381 non-cytotoxic compounds from the five 96-well plates (Plates A to E) of PRB and (B) 204 non-cytotoxic compounds from the three 96-well plates (Plates VEC, MB2 and ZND) of GHPB. The dashed lines indicate the cut-offs for each box. Each dot represents an individual compound, blue dots indicate the 1% DMSO negative control and green dots indicate active compounds considered hits. Images were created with GraphPad Prism 8.0.

To validate the 9 hits identified in the screening assay, dose-response curves were performed in 96-well plates with a two-fold serial dilution of each compound against BUNV-eGFP, with concentrations ranging from 10 µM to 20 nM for determination of the EC_50_ for each compound (**Figure 2**). It was found that three compounds presented the lowest EC_50_ values as demonstrated by Table 1, specifically compounds MMV1580173 (Trimetrexate), MMV690621 (GSK-983) and MMV1634385 inhibited BUNV-eGFP replication by reducing GCU at low concentrations (EC_50_ = 0.5, 2.7 and 1.5 μM, respectively). Selectivity indices (SI) were calculated based on the CC_50_/EC_50_ ratio (**Figure 2**, **Table 1**). For CC_50_ values, due to the limited amount of compound available, concentrations greater than 10 µM could not be tested. Therefore, it was stated that CC_50_ > 10 µM. The three compounds MMV1580173 (Trimetrexate), MMV690621 (GSK-983) and MMV1634385 were taken forward for further analysis.

**Figure 2.**
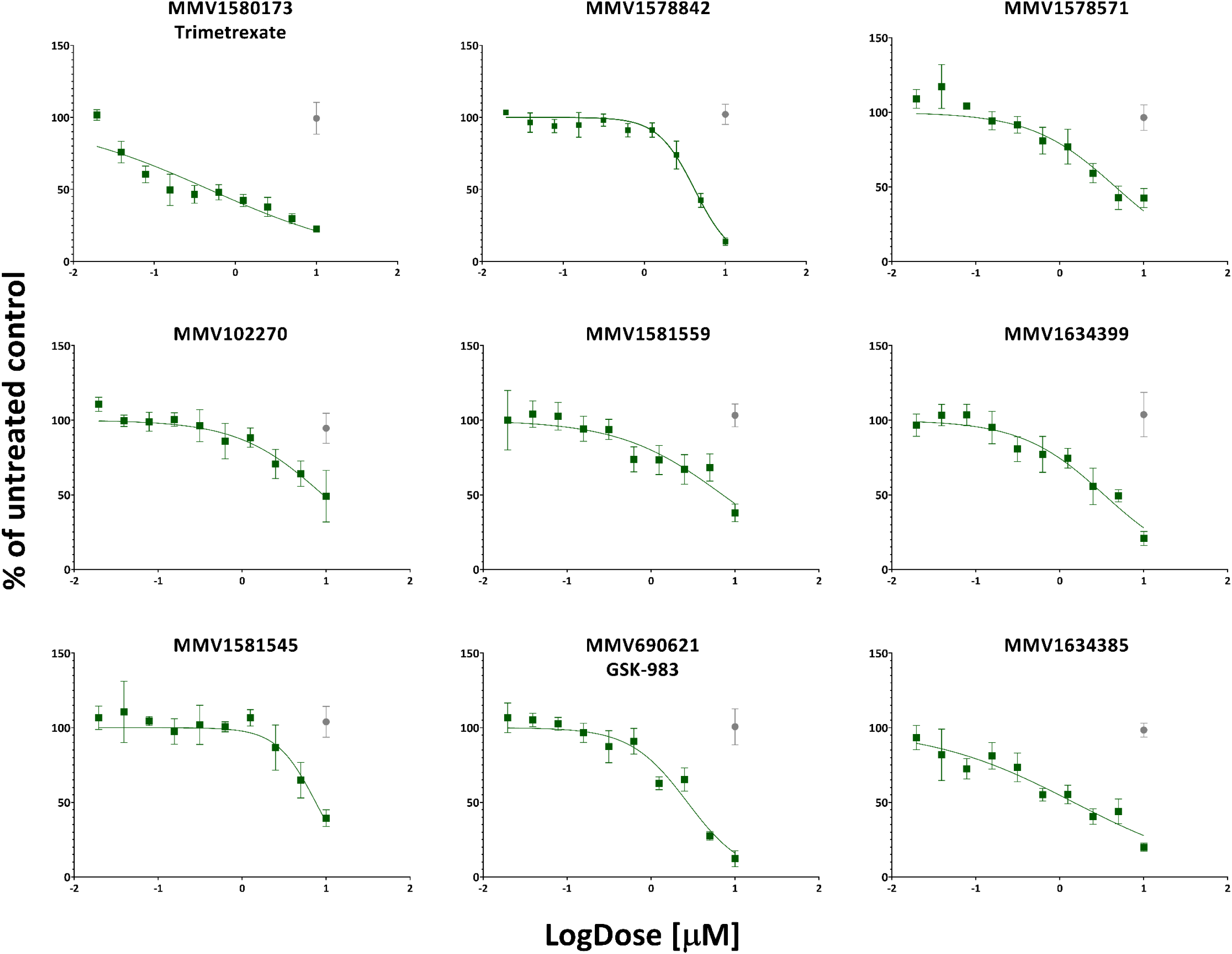
Dose-response (EC_50_) curves of hits from PRB library. A549 cells were incubated with each compound at 2-fold serial dilutions (from 10 µM to 20 nM) for 24 h and eGFP activity was measured in the IncuCyte S3 system (indicated by green lines). Cellular viability at 10 µM is indicated by gray dots. Mean values of three independent experiments each measured in quadruplicate including the standard deviation are shown. Images were created with GraphPad Prism 8.0.

**Table 1:** Antiviral activity of PRB hit compounds against BUNV-eGFP.

| Box | Plate | Compound | Classification | Name | Mean GCU (antiviral assay) | EC<sub>50</sub> (μM) | SI |
| --- | --- | --- | --- | --- | --- | --- | --- |
| PRB | A | MMV1580173 | Antibacterial | Trimetrexate | 155 | 0.5 ± 0.2 | > 20 |
| | B | MMV1578842 | Antibacterial | butyl N-[2-cyclohexyl-6-(diethylamino)-1H-benzimidazol-5-yl]carbamate | 143 | 4.2 ± 0.7 | > 2.4 |
| | | MMV1578571 | Antibacterial | ESI-09 | 146 | 4.6 ± 2.4 | > 2.2 |
| | C | MMV102270 | Antibacterial | Diphyllin | 150 | 9.2 ± 2.6 | > 1.1 |
| | | MMV1581559 | Antibacterial | 5-methyl-3-(3-methylimidazol-4-yl)-7-(2-methylpropyl)-2-(naphthalen-1-ylmethyl)pyrazolo[3,4-d]pyrimidine-4,6-dione | 125 | 7.1 ± 2.1 | > 1.4 |
| | | MMV1634399 | Antibacterial | 4-methyl-8-phenoxy-1-(2-phenylethyl)-2,3-dihydropyrrolo[3,2-c]quinoline | 195 | 3.4 ± 1.5 | > 2.9 |
| | | MMV1581545 | Antibacterial | 2,4,5-trihydroxy-1,3-diiodo-7-methylanthracene-9,10-dione | 123 | 7.5 ± 3.4 | > 1.3 |
| | D | MMV690621 | Antiviral | N-[rac-(1R)-6-chloro-2,3,4,9-tetrahydro-1H-carbazol-1-yl]pyridine-2-carboxamide (rac-GSK-983) | 60 | 2.7 ± 0.7 | > 3.7 |
| | E | MMV1634385 | Antiviral | 4-[1-(1-adamantyl)-5-(trifluoromethyl)benzimidazol-2-yl]aniline (ChEMBL1092703) | 125 | 1.5 ± 0.9 | > 6.7 |

Mean GCU, efficacy (EC<sub>50</sub>) and SI of compounds for inhibition of BUNV-eGFP replication are shown. Mean GCU of each compound was compared with DMSO 1% (mean GCU = 2400) used as vehicle control.

### 3.2. Identified compounds can act in the early and late stages of BUNV replication cycle

To determine which stage of the BUNV replication cycle is affected by the three selected compounds, a time-of-addition experiment was performed. A549 cells were treated with 10 µM of Trimetrexate, GSK-983 or MMV1634385, or an equivalent volume of vehicle control (DMSO). For pre-treatment conditions, cells were exposed to compounds for 1 h prior to infection, after which the compounds were removed before viral inoculation (pre-treatment). For entry assays, compounds were present only during viral adsorption (1 h), after which the inoculum was removed and cells were washed and incubated in compound-free medium for 24 h. To assess a potential virucidal activity, BUNV-eGFP was incubated with each compound for 1 h in the absence of cells, after which the mixture was used to infect A549 cells following the entry assay protocol. For virucidal analyses, a inoculum of compounds and virus was maintaineFor post-infection conditions, compounds were added after viral entry (1 h) and maintained throughout the remainder of the experiment. GCU were quantified at 24 hpi, with additional time points (4, 6, 8, 10, 12, 18 hpi) included for post-infection analyses.

It was shown that none of the three compounds presented effects on pre-treatment or as virucidal agents (**Figure 3A and 3C**). However, GSK-983 and MMV1634385 reduced viral titres by approximately 6-fold when added during BUNV entry assay (p < 0.01), suggesting a possible effect in the early stages of viral infection (**Figure 3B**). Regarding the late-stage analysis, BUNV infection led to a progressive increase in eGFP signals between 6 and 24 hpi in the control group (**Figure 3D**). Treatment with Trimetrexate and GSK-983 significantly reduced GCU at 6 hpi (p < 0.001) when compared to the control, and this effect was maintained up to 24 hpi (p < 0.0001) (**Figure 3D**). In contrast, MMV1634385 did not significantly reduce GCU levels at late time points, indicating a lack of detectable effect on post-entry replication under these conditions.

**Figure 3.**
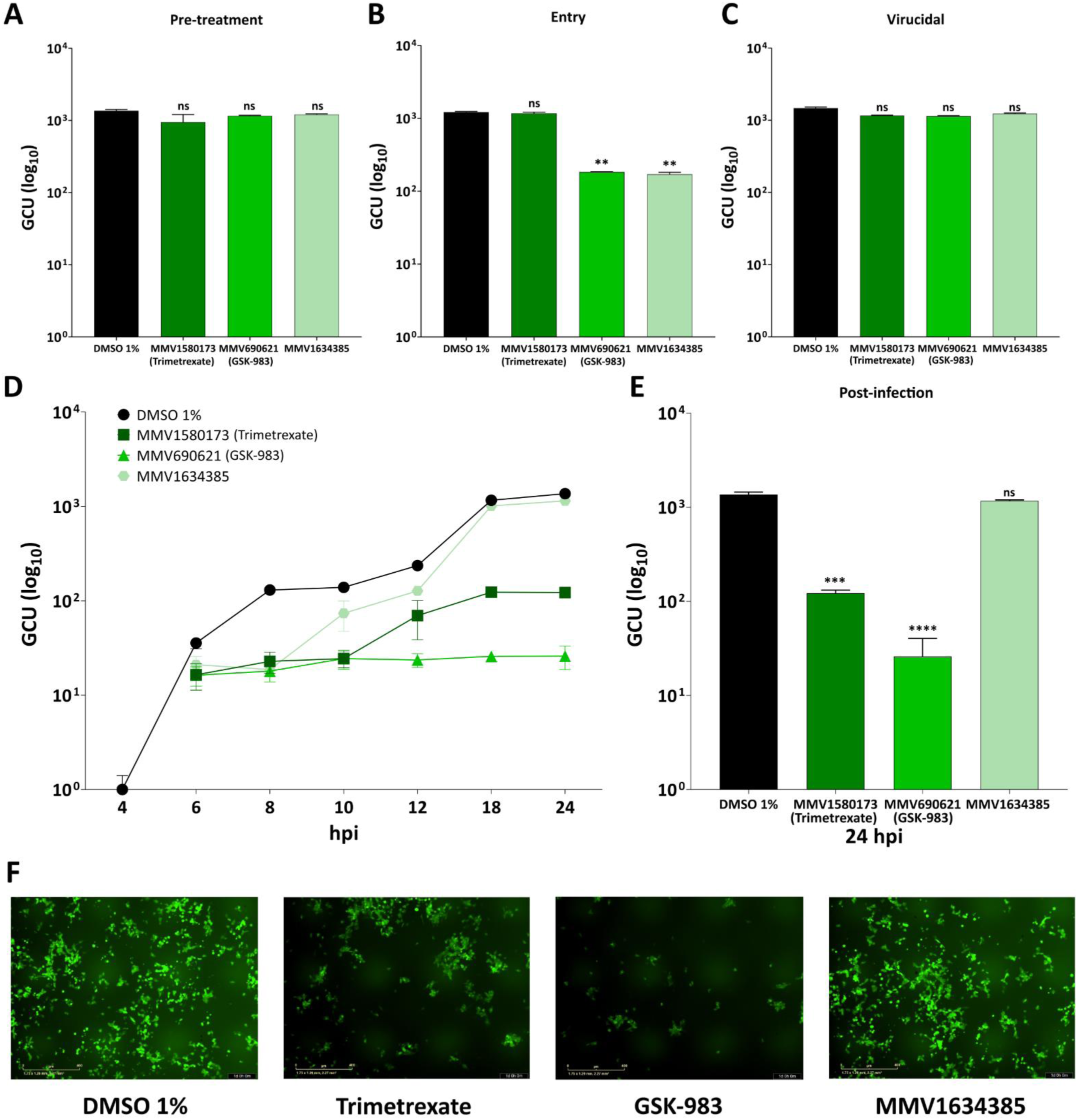
Evaluation of the validated hits on BUNV-eGFP replication cycle. Effects of MMV1580173 (Trimetrexate), MMV6906 21 (GSK-983), and MMV1634385 on cell pretreatment (A), viral entry (B) and as virucidal agents (C) are shown. BUNV-eGFP replication was measured at 24 hpi by GCU enumeration using the IncuCyte S3 system. (D) Time course of eGFP signal (GCU, log_10_) following BUNV infection. Control = black; compounds = shades of green. Compounds were added at 1 hpi; GCU monitored by IncuCyte up to 24 hpi. (E) Quantification of validated hits post-infection treatment at 24 hpi. Values of three independent experiments each measured in quadruplicate including the standard deviation are shown. Images were created with GraphPad Prism 8.0. (F) Representative IncuCyte images at 24 hpi of DMSO 1% control and compounds in the post-infection assay.

To quantify the overall antiviral effect, eGFP intensity at 24 hpi was compared among treatments and confirmed their inhibitory effect of Trimetrexate (11-fold reduction in GCU) and GSK-983 (53-fold reduction), compared to the negative control (p < 0.0001; **Figure 3E and 3F**). Together, these results indicate that Trimetrexate predominantly affects post-entry stages of BUNV infection, GSK-983 impacts both entry and subsequent replication stages, whereas MMV1634385 appears to act primarily during viral entry without a measurable effect on later replication events. The sustained reduction in viral output from 6-8 hpi onwards for Trimetrexate and GSK-983 may reflect inhibition of secondary rounds of viral replication.

### 3.3. Antiviral activity validation of the identified compounds against OROV

To confirm the potential of the identified compounds to inhibit OROV, the new reassortant was used. This strain, named AM0088, was isolated during the recent outbreak of Oropouche fever in Brazil and has significantly higher replicative rates in mammalian cells [12].

The three selected compounds were tested against OROV_AM0088_ in a two-fold serial dilution in 24-well plates. After 5 days, PFU were counted. Results demonstrated that the three hits were able inhibit OROV replication in a dose-dependent manner, as seen for BUNV-eGFP (**Figure 4B**). However, compound GSK-983 was found to have the lowest EC_50_ of 0.5 µM, with a SI > 20, suggesting it as a promising lead compound for further antiviral development against OROV (**Table 2**).

**Figure 4.**
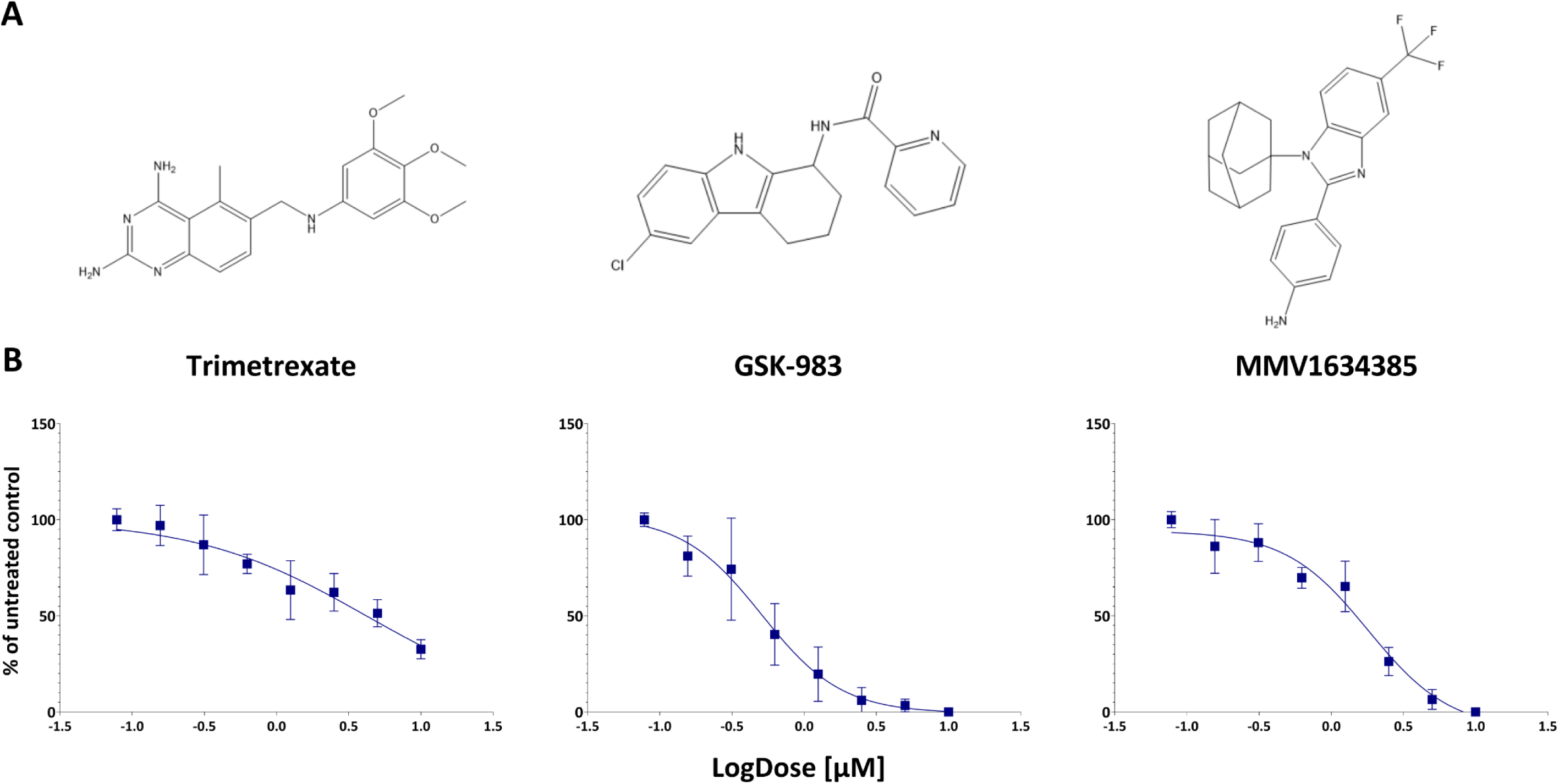
Dose-response (EC_50_) curves of selected compounds against OROV. (A) Chemical structure of Trimetrexate, GSK-983 and MMV1634385 (https://mmv.org/). (B) A549 cells were incubated with each compound at 2-fold serial dilutions (from 10 µM to 20 nM) and PFU were measured after 5 days (indicated by blue lines). Mean values of three independent experiments each measured in triplicate including the standard deviation are shown. Images were created with GraphPad Prism 8.0.

**Table 2.** Antiviral activity of selected compounds against OROV_AM0088_.

| Compound | EC<sub>50</sub> (μM) | SI |
| --- | --- | --- |
| Trimetrexate | 4.2 | > 2.4 |
| GSK-983 | 0.5 | > 20 |
| MMV1634385 | 1.8 | > 5.6 |

## 4. Discussion

According to the World Health Organization (WHO), Brazil is one of the countries with the highest number of arboviruses cases [24]. Extensive natural ecosystems, tropical climate and densely populated urban areas provide ideal conditions for their transmission and spread [25]. The re-emergence of arboviruses such as OROV, in addition to the lack of specific antivirals for many of these pathogens, highlights the urgency of innovative approaches to combat outbreaks and reduce the public health impact of infections [26]. Here, we developed a medium-throughput screening assay with an automated image-based readout using a system of BUNV-eGFP as a model to identify antiviral drugs against OROV.

Since the drug discovery for emerging or re-emerging viruses has currently been accelerated by the exploration of new molecular pathways and intervention targets of well-characterized drugs, whether approved or under investigation, open-source libraries aim to identify new treatments for viral infections beyond their original indication, characterizing the so-called drug repositioning [13]. Considering that they have already demonstrated safety in humans, repositioned drugs can not only reduce the failure rate in developing a new antiviral candidate, but also minimize the time and resources required for the discovery of a new drug [27]. MMV libraries provide a set of structurally diverse compounds for screening against infectious and neglected diseases. The compounds selected from a comprehensive list of antibacterial, antiviral, and antifungal agents, have diverse mechanisms of action. Many of them are already on the market or in stages of drug discovery and development. Their biological activity and selection were based on information available in the literature, saving time and resources by establishing a foundation for the subsequent development of therapeutic approaches [28,29].

Here, our screening assay allows for the precise and accurate screening of these libraries, and the rapid measurement of BUNV titres. Furthermore, its large-scale range makes it highly suitable for high-throughput screening of potential antiviral compounds against emerging peribunyaviruses. With the identification of 9 hits (1.5%), being 7 antibacterials and 2 antivirals, our data showed that 3 of them (Trimetrexate, GSK-983 and MMV1634385) exhibit lower EC_50_ values and consequently higher SIs. Despite the limitations in identifying the exact CC_50_s due to compound availability, the absence of cytotoxicity up to 10 µM provides a reasonable indication of a favourable safety margin at this stage of evaluation.

Compound MMV1580173, known as Trimetrexate, exhibited a low EC_50_ against BUNV-eGFP and for this reason was selected to be validated against OROV. It is a folinic acid analogue belonging to the same drug class of methotrexate (MTX), a widely used chemotherapy and immunosuppressant drug [30,31]. Trimetrexate exerts its action by inhibiting dihydrofolate reductase (DHFR), an enzyme related to the de novo synthesis of nucleosides needed for nucleic acid production [32]. It has been used as a treatment for infection of *Pneumocystis carinii*, reducing the production of DNA and RNA precursors and acting as an anti-inflammatory due to subsequent cellular effects [33]. These activities have been related as beneficial to many repurposing approaches, including as an antiviral [34,35]. Iaconis and coworkers described Trimetrexate as an inhibitor of SARS-CoV-2 entry on Vero E6 and A549 cells. Furthermore, the influence of this drug on the viral nsp13 was identified, highlighting its polypharmacological profile and corroborating our post-infection results [36]. Despite not displaying the most favorable EC_50_ and SI values against OROV, Trimetrexate still demonstrated significant inhibitory effects, suggesting its potential as a candidate for further study.

Compound MMV1634385 (CHEMBL1092703) is a benzimidazole derivative featuring adamantyl and trifluoromethyl substituents [37]. While specific studies on its exact structure are limited, related compounds in the benzimidazole class have demonstrated notable biological activities, suggesting potential properties for this compound [37]. Benzimidazole derivatives have been reported to exhibit antiviral activity against many RNA and DNA viruses, including Coxsackie virus B3 (CSV), hepatitis C virus (HCV), respiratory syncytial virus (RSV), Herpes simplex 1 virus (HSV-1) and yellow fever virus (YFV) [38,39]. Their effects include inhibition of genome replication and protein processing as evidenced by the impairment of the HCV NS5B polymerase [39]. Notably, in our results, MMV1634385 did not exhibit a detectable inhibitory effect on post-entry BUNV replication, suggesting that its antiviral activity may differ from that reported for other benzimidazole derivatives and viral families. Instead, our data indicate an effect restricted to early stages of infection. Such divergence may reflect virus-specific replication strategies, differences in compound-virus interactions or structural properties of MMV1634385 [38]. As described by Wilkinson and coworkers (2017), some derivatives can interfere with early stages of viral replication, such as viral entry into host cells [40]. Together, these observations highlight the mechanistic diversity of benzimidazole derivatives across different viral families.

The most notable compound identified was MMV690621, a tetrahydrocarbazole inhibitor of the dihydroorotate dehydrogenase (DHODH), an antiviral drug named GSK-983, with a broad-spectrum activity via a host target [41]. Although it did not display the greatest EC_50_ and SI values against BUNV, this compound showed the strongest inhibitory activity in the screening assay, a significant constant inhibition of BUNV replication, and exhibited the best performance against OROV. It has been described as a drug that can inhibit a diversity of viruses, from different families, with EC_50_s values varying from 5-20 nM [41]. Our research group had previously characterized its activity against chikungunya virus (CHIKV), where it displayed an EC_50_ of 13.8 µM and a CC_50_ > 100 µM in BHK-21 cells [42]. Here, an EC_50_ of 2.7 µM was determined against BUNV, and of 0.5 µM against OROV. An exact SI could not be calculated, however, the lack of cytotoxicity at the highest concentration tested suggests that this compound has a highly favorable selectivity profile, which still reflects its promising safety and efficacy profile against these orthobunyaviruses. As a drug that functions through a cellular target, it is expected that it can interfere with multiple stages of the viral replicative cycle and that it can act against many viral families. The inhibition of DHODH depletes the host cell pyrimidine content, reducing intracellular UTP and CTP levels, and consequently limiting nucleoside triphosphates available for viral genome replication and transcription [43]. In addition, it can impair glycosylation and other pyrimidine-dependent processes in host cells, further stressing viral replication [44]. To best of our knowledge, this is the first report of GSK-983 acting as an anti-orthobunyavirus compound, expanding the antiviral spectrum of this drug.

In conclusion, our results demonstrated that our screening assay was able to screen and identify antiviral compounds against BUNV-eGFP in the PRB library from MMV. 3 compounds were identified as hits and GSK-983 demonstrated to have the greatest effects in the validation assays. Future studies should evaluate the antiviral efficacy of GSK-983 and assess its pharmacokinetic profile. Additionally, the BUNV-eGFP system may be adapted for high-throughput screening of other *Orthobunyavirus* members, providing a scalable tool for rapid antiviral discovery.

## Acknowledgments

We would like to thank the Medicines for Malaria Venture (MMV, www.mmv.org) for the design and supply of the Pandemic Response Box and Global Health Priority Box. We also would like to thank Professor José Proença Modena for the provision of OROV_AM0088_. ACGJ is grateful for funding from to FAPEMIG (APQ-01487-22 and APQ-04686-22) and to CAPES (Prevention and Combat of Outbreaks, Endemics, Epidemics and Pandemics— Finance code #88881.506794/2020-01 and Finance code #001).

## Data availability statement

All data generated or analysed during this study are included in this published article (and its Supplementary material).

## Author contributions

NMC, MS and ACGJ conceptualized the study. NMC and HMP performed the investigation, analysis and data curation. NMC wrote the original draft. HMP, JNB, MS, and ACGJ performed subsequent manuscript reviewing and editing. JNB, MS and ACGJ involved in supervision. MS and ACGJ were responsible for funding acquisition.

