## Supplementary material for "Screening of MMV open-source libraries using Bunyamwera virus as a model reveals inhibitors of Oropouche virus infection"

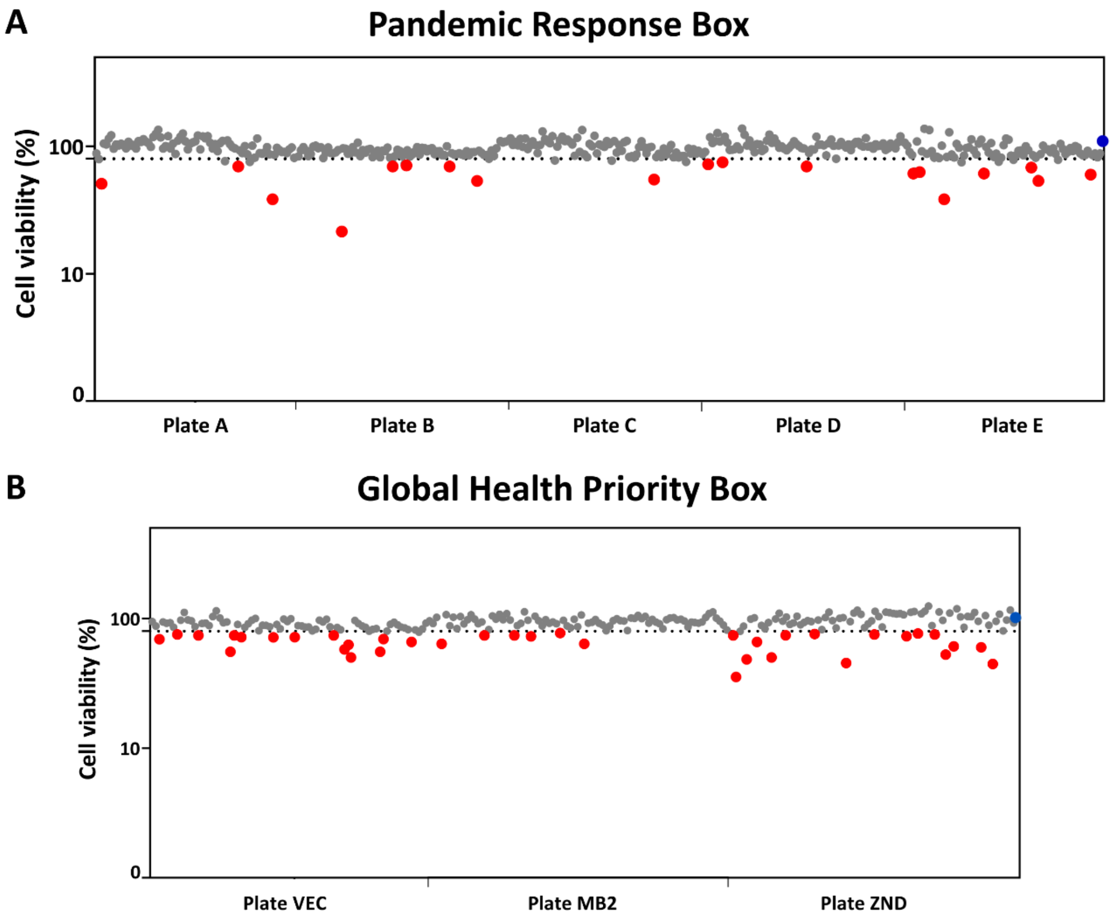


**Figure S1. Cytotoxicity of MMV boxes on A549 cells.** The dashed lines indicate the cut-offs (< 80%). Each dot represents an individual compound, blue dots indicate the 1% DMSO vehicle control and red dots indicate cytotoxic compounds. Images were created with GraphPad Prism 8.0.
